# Temporal dynamics of peri-microsaccadic modulations within the foveola

**DOI:** 10.1101/2025.10.29.685430

**Authors:** Zoe Stearns, Martina Poletti

**Author notes:** **For correspondence:** (MP). Meliora Hall, University of Rochester, Rochester, NY, USA, 14627.

## Abstract

Microsaccades are small, rapid eye movements that shift the center *of* gaze by less than half a degree. While they have traditionally been associated with perceptual modulations and covert attention shifts in extrafoveal vision, recent evidence indicates that microsaccades also modulate perception within the central fovea in a spatially selective manner. However, the temporal dynamics of these modulations in fine spatial discrimination remain largely unexplored. Here we used high-precision eye tracking and gaze-contingent display control to achieve accurate localization of the line of sight and to restrict visual stimulation to selective locations within the central fovea during instructed microsaccades. Our results show that visual performance peaks approximately 70 milliseconds before microsaccade onset at the target location, while perception at equally eccentric, non-target foveal locations is concurrently impaired. This is followed by a generalized suppression phase while the microsaccade is in flight, when fine spatial discrimination drops to near-chance levels. Visual performance then recovers rapidly at the target location, returning to baseline within 100 milliseconds after microsaccade landing. Our findings demonstrate that visual discrimination within the central fovea is modulated following a distinct temporal profile and in a spatially selective way around the time of microsaccade execution.

## Introduction

Humans sample the visual scene with saccades, bringing objects of interest into the central fovea for finer examination. However, even during fixation, small microsaccades, tiny saccades less than half a degree in size, continually occur (***Rolfs, 2009***; ***Kowler, 2011***; ***Poletti and Rucci, 2015***). Microsaccades are ubiquitous in natural oculomotor behavior. Present in various activities, from reading to fine visuomotor tasks (***Bowers and Poletti, 2017***; ***Poletti et al., 2013***; ***Ko et al., 2010***), these tiny movements can occur 0.5 to 2 times per second. Although microsaccades were initially thought to prevent image fading (***Collewijn and Kowler, 2008***; ***Martinez-Conde et al., 2004***) and correct for fixational offsets introduced by ocular drift (***Cornsweet, 1956***; ***Engbert and Mergenthaler, 2006***), there is now mounting evidence showing that these tiny gaze shifts are finely controlled (***Ko et al., 2010***; ***Shelchkova and Poletti, 2020***; ***Poletti et al., 2020***) and enable active exploration of complex foveal scenes (***Shelchkova et al., 2019***; ***Intoy et al., 2021***). In addition, they introduce temporal transients that reshape the power of visual input in a unique way (***Mostofi et al., 2020***). Microsaccades have also gained popularity in recent years as they appear to act as an index of peripheral covert attention (***Yuval-Greenberg et al., 2014***; ***Yu et al., 2022***; ***Liu et al., 2024***; ***Gu et al., 2024***). While microsaccades can be used to index attention in conditions in which subjects are required to maintain fixation on a point while peripheral stimuli are presented or cued, normally, when the foveal stimulus is rich in detail, microsaccades are engaged in visual exploration. Although most of the research has focused on the link between microsaccades and covert attention in the visual periphery, the interplay between microsaccades and fine spatial attention in the foveola has been less examined. A better understanding of this interplay is important to determine how foveal vision is modulated when humans engage in visually guided fine motor skills, examine fine spatial patterns, and when they read.

Previous work by (***Shelchkova and Poletti, 2020***) has shown that the preparation of microsaccades leads to perceptual modulations in foveal vision. Specifically, perceptual performance at the microsaccade target location is enhanced during the preparation period, whereas non-target locations are perceptually impaired. (***Guzhang et al., 2024***) further examined the spatial resolution of these pre-microsaccadic perceptual modulations in the foveola, finding that the enhancement at the microsaccade goal location is highly localized in a region of 10’ radius around the microsaccade target. These findings indicate that microsaccades and their preceding attentional shifts reshape visual perception across the central fovea in a spatially selective way. However, the temporal dynamics of these pre-microsaccadic modulations remain unknown. How long before the microsaccade begins is vision enhanced at the microsaccade goal location? In both studies, the target stimuli were shown around 100 milliseconds after the motor cue onset. This consistent timing ensured pre-microsaccadic exposure of the stimuli, but did not allow to resolve at a fine temporal scale the peri-microsaccadic changes of visual perception. Further, in both studies target duration was too long to capture the rapid perceptual dynamics unfolding before microsaccade onset.

Recent work by (***Intoy et al., 2021***) observed dynamic and non-uniform attenuations of foveal contrast sensitivity during the execution of microsaccades, showing that the region closest to the center of gaze undergoes the greatest and most rapid modulation compared to more eccentric foveal regions. Consistent with (***Shelchkova and Poletti, 2020***; ***Guzhang et al., 2024***), (***Intoy et al., 2021***) report an improvement in visual detection near the microsaccade target location before microsaccade onset with respect to non-target locations at comparable eccentricity. However, it is not clear when this modulation begins with respect to the microsaccade onset, for how long it lasts, and whether it is the result of a suppression at the non-foveated location, an enhancement at the microsaccade target, or both. Further, (***Intoy et al., 2021***) examined contrast sensitivity using a spot detection task that did not require fine spatial discrimination. Therefore it is still unclear how perimicrosaccadic fine spatial vision is temporally modulated. This study aims to fill this gap and to measure the time course of pre-microsaccadic attention and peri-microsaccadic modulations of fine spatial vision.

Examining the time-course of perception at spatially selected locations within the 1-deg central fovea is technically challenging. Microsaccades can be as small as a quarter of a degree (***Poletti et al., 2020***) and last 30-80 milliseconds, requiring precise and accurate measurement and monitoring of gaze position during fixation. The perceptual enhancement preceding each microsaccade execution is highly localized (***Guzhang et al., 2024***), meaning both the starting and landing positions of the microsaccade must be rigorously monitored in relation to the stimulus position. Consequently, examining fine spatial discrimination around the microsaccade goal location on a fine-grained temporal scale requires not only high-precision eye-tracking apparatus but also accurate localization of the line of sight. The interpretation of eye movement behavior can be significantly complicated by various factors, including head movements, dynamic pupil size changes, and corneal reflections. These elements can confound eye position signals, making it challenging to accurately assess perceptual effects (***Morimoto and Mimica, 2005***; ***Wyatt, 2010***; ***Morgante et al., 2012***; ***Choe et al., 2016***). To circumvent these problems, we used a high-precision Dual Purkinje Image (DPI) eye-tracker coupled with a system for gaze-contingent display control capable of localizing the line of sight with arcminute precision. This setup allowed us to examine the perceptual changes in foveal vision that begin during the microsaccade preparation period and extend throughout the relocation of the center of gaze. Our results reveal that pre-microsaccadic modulations within the foveola begin to unfold ≈100 milliseconds before the microsaccade onset, peak ≈70 milliseconds before, and end ≈30 milliseconds prior to the movement onset. We report a rapid recover from pre-microsaccadic suppression upon landing. These rapid pre-microsaccadic changes were not reported when subjects failed or were not instructed to execute microsaccade.

## Results

To explore the temporal dynamics of peri-microsaccadic perceptual modulations in foveal vision, we assessed observers’ ability to discriminate the orientation of high-acuity stimuli located a few arcminutes away from the central fixation marker at various time points relative to microsaccade onset. Each trial started with observers fixating on the central marker. In two-thirds of the trials, a motor (directional) cue prompted observers to rapidly shift their gaze to the cued location (Figure 1C, left), while on the remaining trials, a neutral cue instructed observers to maintain fixation at the central position (Figure 1C, right). Following the onset of either cue, two oriented bars were briefly presented after varying delays. Positioned ±18 arcminutes away from the central fixation marker, both stimuli fell well within the central one-degree foveola. A response cue, a vertical line above one of the stimuli locations, appeared 500 milliseconds after stimulus presentation. Observers were asked to report the orientation (±45 deg) of the tilted bar stimulus presented at the location indicated by the response cue. In motor cue trials, the response cue could be spatially congruent or incongruent with the motor cue prompting the microsaccade (Figure 1B). As illustrated in Figure 1D, stimuli were presented at comparable delay times from the trial onset in both motor and neutral cue trials (average and standard deviation across observers: 391±10ms vs 377±19ms, *p*=0.06; paired-sample t-test), particularly in trials corresponding to the pre-microsaccadic interval (average and standard deviation across observers: 248±19ms vs 239±22, *p*=0.09; paired-sample t-test. See Methods: *Calculating Baseline Performance in Neutral Trials* under the *Performance Analysis* section for how delay times relevant to pre-microsaccadic interval are determined for neutral trials). Therefore, spatially and temporally, retinal stimulation was equivalent in neutral and motor trials. In motor cue trial, stimuli onset relative to microsaccade onset was uniformly distributed in the period -150 milliseconds to + 300 milliseconds after microsaccade onset(Figure 1E). Eye movements in the task were recorded using a high-precision Digital Dual Purkinje Image eye tracker, enabling precise localization of the line of sight during high-acuity tasks (***Santini et al., 2007***; ***Wu et al., 2023***; ***Ko et al., 2010***).

**Figure 1.**
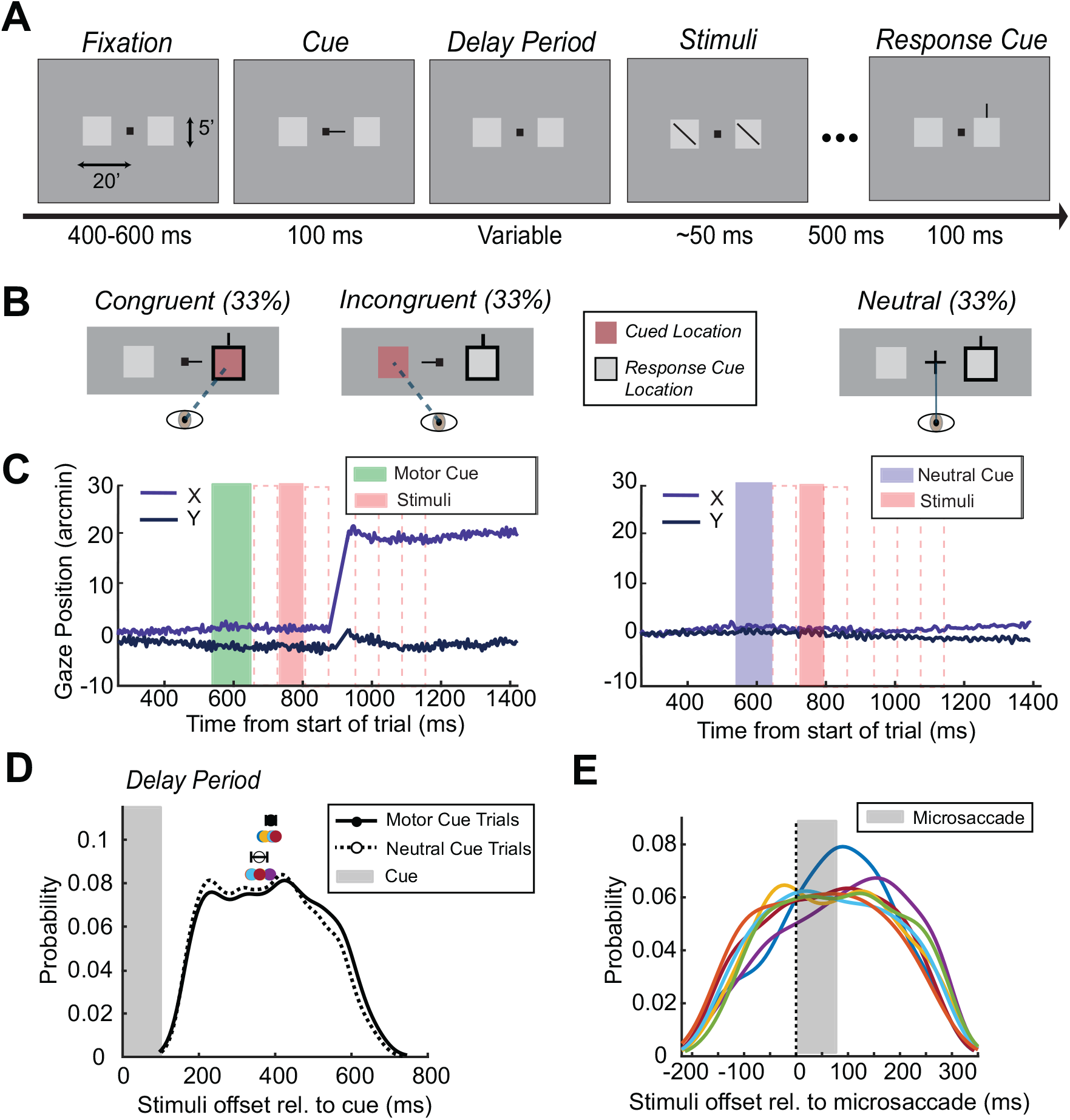
Experimental Paradigm. *A*) Observers began each trial by fixating on a central dot surrounded by two squares (5’ x 5’ in size) each offset 18 arcmin to the left/right of the center. A central cue instructed observers to either shift their gaze to the left/right location or maintain fixation at the center (neutral condition). After a variable delay period two fine spatial stimuli, bars tilted ±45deg, were briefly (50 milliseconds) presented at both locations. After 500 milliseconds a response cue appeared above either the left or right location. Observers were instructed to report the orientation of the stimulus previously shown at the location highlighted by this cue. *B*) The motor cue could be either spatially congruent or incongruent with the response cue location. In the neutral trials a fixation cross instructed subjects to maintain the gaze at the center. *C*) Examples of gaze position during a typical motor (Left) and neutral (Right) trial. Dashed vertical lines indicate stimuli presentation. *D*) Probability of stimuli presentation time relative to the motor cue (solid line) and neutral cue (dashed lines) trials. *E*) Probability of stimuli presentation relative to the microsaccade onset. Dashed line marks microsaccades onset time. Probability distributions in *D*) and *E*) were calculated based on a nonparametric kernel-smoothing. Individual observers are color-coded.

Consistent with previous work (***Shelchkova and Poletti, 2020***), a directional motor cue presented at central fixation effectively guides overt attention at the foveal scale. As depicted in Figure 2, observers executed precise microsaccades directed towards the location indicated by the endogenous motor cue. The average amplitude of the microsaccade was measured at 19 ± 6 arcminutes, approximately matching the spatial offset of the stimulus from the fixated marker (Figure 2A). The average 2-D landing error, defined as the distance between the microsaccade landing position and the center of the target stimulus, was 7.5±1.4 (Figure 2C). Not only were subjects’ microsaccades accurate, but they were also executed in a timely manner; the average microsaccade latency, the time between cue onset and microsaccade onset, was 326 ± 34 milliseconds (Figure 2B). Microsaccade latency values reported here are also consistent with previous reports for microsaccades of similar amplitude using similar experimental paradigms (see (***Shelchkova and Poletti, 2020***; ***Guzhang et al., 2024***)).

**Figure 2.**
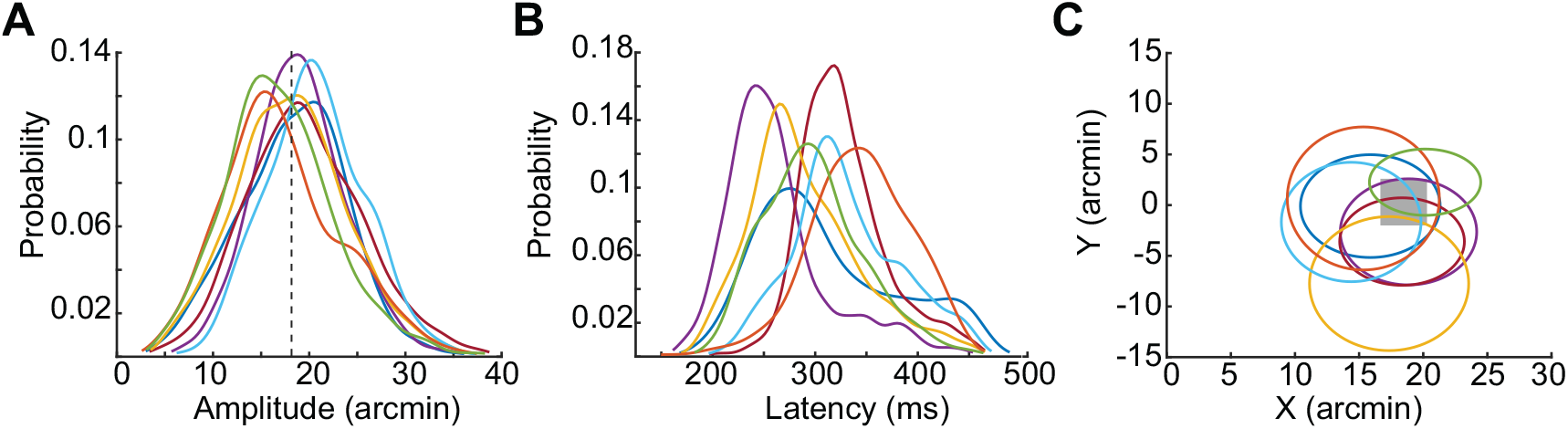
Microsaccade characteristics. Microsaccade amplitude (*A*) and latency (*B*) distributions. Dashed lines in (*A*) show the target distance from the center of the display. *C*) 68% confidence ellipses of microsaccade landing endpoints. The gray square represents the location of the target drawn to scale. Probability distributions in *A*) and *B*) were calculated based on a nonparametric kernel-smoothing. Individual observers are color-coded.

### Visual Discrimination Before Microsaccades

We first examined the modulation of fine visual discrimination before the time of instructed microsaccades. As depicted in Figure 3B, visual discrimination (measured as *d*^′^) underwent a significant modulation in the pre-microsaccadic interval, during which gaze remained at the center (see Figure 3A, Before). Approximately 100 to 150 milliseconds before the onset of the microsaccade, performance in congruent and incongruent trials was comparable to baseline (neutral) and above chance level (*d*^′^= 1.17±0.31, *d*^′^= 1.43±0.88, *d*^′^= 0.85±0.48 for neutral, congruent and incongruent respectively; see Supplementary Figure 6 one-way repeated-measures ANOVA, condition:*F*(2,12)= 1.25; *p*= 0.32). However, ≈100 milliseconds before the onset of the microsaccade, visual discrimination progressively increased in congruent trials, while at the same time decreasing in incongruent trials (Figure 3C; one-way repeated-measures ANOVA, condition:*F*(2,12)= 21.21; *p*= 0.0001; Tukey-Kramer corrected *t*-tests: *p*= 0.0665, *BF*_10_= 2.93, Cohen’s *d*= 1.07 for neutral before vs. congruent before; *p*= 0.0320, *BF*_10_= 5.23, Cohen’s *d*= -1.30 for neutral before vs. incongruent before; *p*= 0.0009, *BF*_10_= 89.21, Cohen’s *d*= -2.70 for congruent before vs. incongruent before). While the enhancement at the target location did not reach significance relative to baseline, the impairment at the non-target location did, and visual discrimination differed substantially between the two locations prior to microsaccade onset.

**Figure 3.**
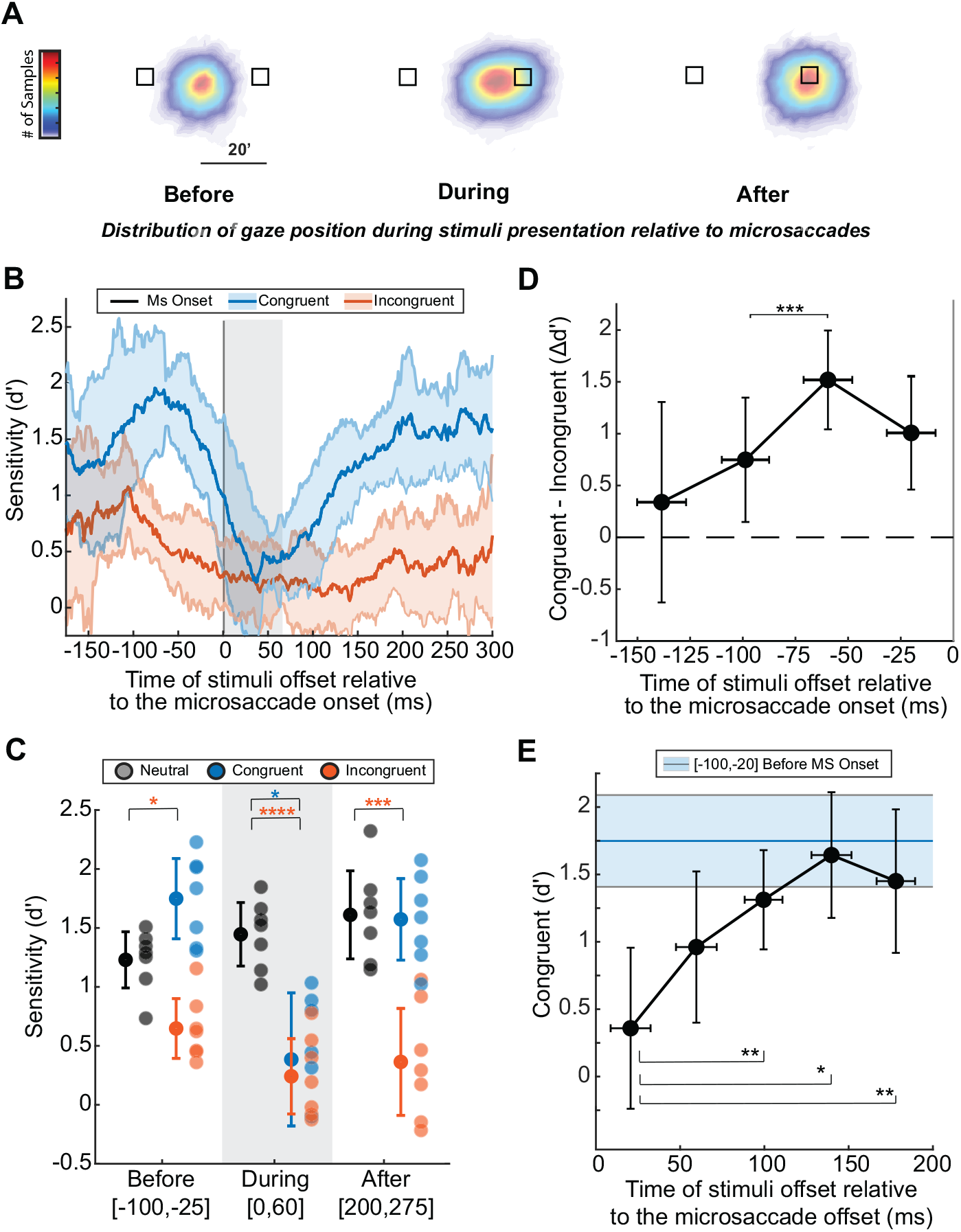
Temporal dynamics of peri-microsaccadic visual discrimination within the foveola. *A*) 2-D probability distribution of gaze position during the stimuli presentation before, during, or after the microsaccade. *B*) Average performance measured as *d*^′^ smoothed as a function of time (window = ±24 ms, step = 1 ms) relative to the microsaccade onset in the congruent (blue) and incongruent (orange) conditions. Shaded error bars represent the margin of error for a 95% within-subject confidence interval for each time bin. Shaded grey region shows microsaccade approximate duration. *C*) Average performance at different times with respect to the microsaccade onset and in delay-matched neutral trials. Shaded dots represents single subject data. ^*^*p<*0.05, ^***^*p<*0.001, ^****^*p<*0.0001 (one-way repeated-measures ANOVA (factor: condition) per each time frame (before, during and after), Tukey–Kramer corrected *t*-tests; comparisons relative to neutral trials are shown). (*D*) Average difference in *d*^′^ between the congruent and incongruent trials at different time points before the onset of the microsaccade. ^*^*p<*0.05, ^**^*p<*0.01, ^***^*p<*0.001 (one-way repeated-measures ANOVA (factor: time) with Tukey-Kramer corrected *t*-tests). (*E*) Average performance in congruent condition at different time points after the offset of the microsaccade. ^*^*p<*0.05, ^**^*p<*0.01 (one-way repeated-measures ANOVA (factor: time) with Tukey-Kramer corrected *t*-tests). Vertical error bars in *C*), *D*), and *E*) are 95% within-subject confidence intervals and horizontal error bars in *D*) and *E*) represent standard deviations.

The average difference in performance between congruent and incongruent trials (Δ*d*^′^) during the pre-microsaccadic interval was 1.05±0.46 *d*^′^, reaching a maximum of 1.52±0.52 approximately 60 milliseconds before the start of the microsaccade (shown in Figure 3D; one-way repeated-measures ANOVA, time:*F*(3,18)= 5.9, *p*= 0.0055; Tukey-Kramer corrected *t*-tests: *p*= 0.06, *p*= 0.0009, *p*= 0.317 for the 3rd time bin compared to the 1st, 2nd, and 4th, respectively). These results show that approximately 100 milliseconds before microsaccade onset, discrimination rapidly improved at the intended target location while decreasing at the non-target location. During microsaccade preparation, fine spatial vision at isoeccentric locations was dynamically and differentially modulated.

### Visual Discrimination During Microsaccades

Previous studies have reported impaired detection performance (***Zuber and Stark, 1966***) and reduced contrast sensitivity (***Intoy et al., 2021***) within the central fovea when stimuli are presented during microsaccades. Consistent with these findings, our results show decreased fine visual discrimination in both congruent and incongruent trials during microsaccade execution (Figure 3C; one-way repeated-measures ANOVA, condition:*F*(2,12)= 13.446; *p*= 0.0009; Tukey-Kramer corrected *t*-tests: *p*= 0.025,*BF*_10_= 6.35, Cohen’s *d*= -1.378 for neutral during vs. congruent during; *p*= 0.00008, *BF*_10_= 666.49, Cohen’s *d*= -4.197 for neutral during vs. incongruent during; *p*= 0.892, *BF*_10_= 0.39, Cohen’s *d*= 0.174 for congruent during vs. incongruent during).

Visual suppression occurs at target and non-target locations and begins slightly before microsaccade onset, in line with prior reports for microsaccades (***Intoy et al., 2021***; ***Zuber and Stark, 1966***) and saccades (***Diamond et al., 2000***; ***Crevecoeur and Kording, 2017***). Our results reveal that fine spatial vision at the target location begins to sharply decline in the final 25 milliseconds leading up to microsaccade onset.

### Visual Discrimination After Microsaccades

We then examined how visual performance across the foveola is affected immediately after microsaccade landing. Notably, in congruent trials, the stimulus is no longer positioned 18 arcminutes away from the preferred locus of fixation (PLF), rather, once the microsaccade lands it appears at the PLF; within approximately 200 milliseconds after microsaccade offset, gaze distance from the target was 8’±2’. We assessed how quickly performance recovered at the PLF after microsaccade suppression. Our findings reveal a rapid increase in performance from 0.36±0.65 to 1.31±0.40, a 0.95±0.36 d-prime difference, within 100 milliseconds after microsaccade offset (Figure 3E; one-way repeated-measures ANOVA, time:*F*(4,24)= 11.04, *p*= 0.00003; Tukey-Kramer corrected *t*-tests between the 1st vs. 2nd, 3rd, 4th, and 5th time bin respectively: *p*= 0.2343, *BF*_10_= 1.892, Cohen’s *d*= 0.907; *p*= 0.0025, *BF*_10_= 81.119, Cohen’s *d*= 2.643; *p*= 0.0293, *BF*_10_= 10.843, Cohen’s *d*= 1.606; *p*= 0.0020, *BF*_10_= 98.944, Cohen’s *d*= 2.766). Eye speed was stable immediately following microsaccade offset and later on (t-test, two-sided, paired: 0.69±0.10^◦^∕*s* vs. 0.73±0.11^◦^∕*s* for eye speed 0–100 ms vs. 200–300 ms post-landing; *p*= 0.49), suggesting that the observed perceptual improvement was unlikely to be driven by retinal factors.

Beyond this rapid post-microsaccadic recovery, performance at the PLF was comparable to that 25-100 milliseconds before microsaccade onset, when the target was 18 arcminutes away (Bonferroni corrected t-test, two-sided, paired: *p*= 0.89, *BF*_10_= 0.461, Cohen’s *d*= -0.309 for congruent before vs. congruent after). It was also improved relative to neutral trials with matched premicrosaccadic delay times (Bonferroni corrected t-test, two-sided, paired: *p*= 0.0094, *BF*_10_= 16.54, Cohen’s *d*= 1.797 for neutral before vs. congruent after), where the stimuli in baseline trials were presented 239±22 milliseconds after cue onset (for more information on how pre-microsaccadic delay times were determined for neutral trials see Methods: *Calculating Baseline Performance in Neutral Trials*).

On the other hand, performance in incongruent trials was impaired relative to baseline after microsaccade landing (one-way repeated-measures ANOVA, condition:*F*(2,12)= 30.259; *p*= 0.00002; Tukey-Kramer corrected *t*-tests: *p*= 0.00033, *BF*_10_= 205.604, Cohen’s *d*= -3.257 for neutral after vs. incongruent after; *p*= 0.972, *BF*_10_= 0.361, Cohen’s *d*= -0.086 for neutral after vs. congruent after; *p*= 0.0042, *BF*_10_= 26.41, Cohen’s *d*= 2.024 for congruent after vs. incongruent after). The persistently poor performance in incongruent trials after the microsaccade is likely attributable to the fact that the cued stimulus is now 35’±2’ away from the PLF (Figure 3A) where fine discrimination is worse (***Poletti et al., 2013***) and contrast sensitivity is reduced relative to the center of gaze (***Intoy et al., 2021***).

### Pre-Microsaccadic Visual Discrimination Relative to the Motor Cue

Thus far, our results have shown a perceptual advantage at the microsaccade target location relative to an isoeccentric location in fine spatial vision beginning 100 milliseconds before microsaccade onset (Figure 4A; two-way repeated-measures ANOVA, condition:*F*(1,6)= 18.13, *p*= 0.00534; time:*F*(3,18)= 2.037, *p*_GG_= 0.168; interaction:*F*(3,18)= 5.902, *p*_GG_= 0.025; Tukey-Kramer corrected *t*-tests: *p*= 0.424, *p*= 0.0022, *p*= 0.000235, *p*= 0.00407 for congruent vs. incongruent at 1st, 2nd, 3rd, and 4th time bin respectively). Yet, it is possible that this pre-microsaccadic effect is influenced by endogenous attention, given that covert attention can be voluntarily controlled at this fine scale (***Poletti et al., 2017***). Voluntary spatial attention is typically manipulated with centrally presented directional cues that indicate likely target locations (***Posner, 1980***; ***Bowman et al., 1993***; ***Van der Stigchel and Theeuwes, 2007***). This classic approach incentivizes the allocation of attention at one spatial location over others, leading to enhanced discrimination at selected locations in the absence of gaze shifts. In our study, the motor cue and the response cue were independent, *i*.e., one did not predict the other (cue validity was 50%). Therefore, since each location was equally likely to be cued for report, there was little incentive to prioritize either location over the other, dampening the benefit of directed attention to a single location on a given trial. However, the delays between the central cue and target onset were within the time period during which the effects of endogenous covert attention usually arise (***Carrasco, 2011***).

**Figure 4.**
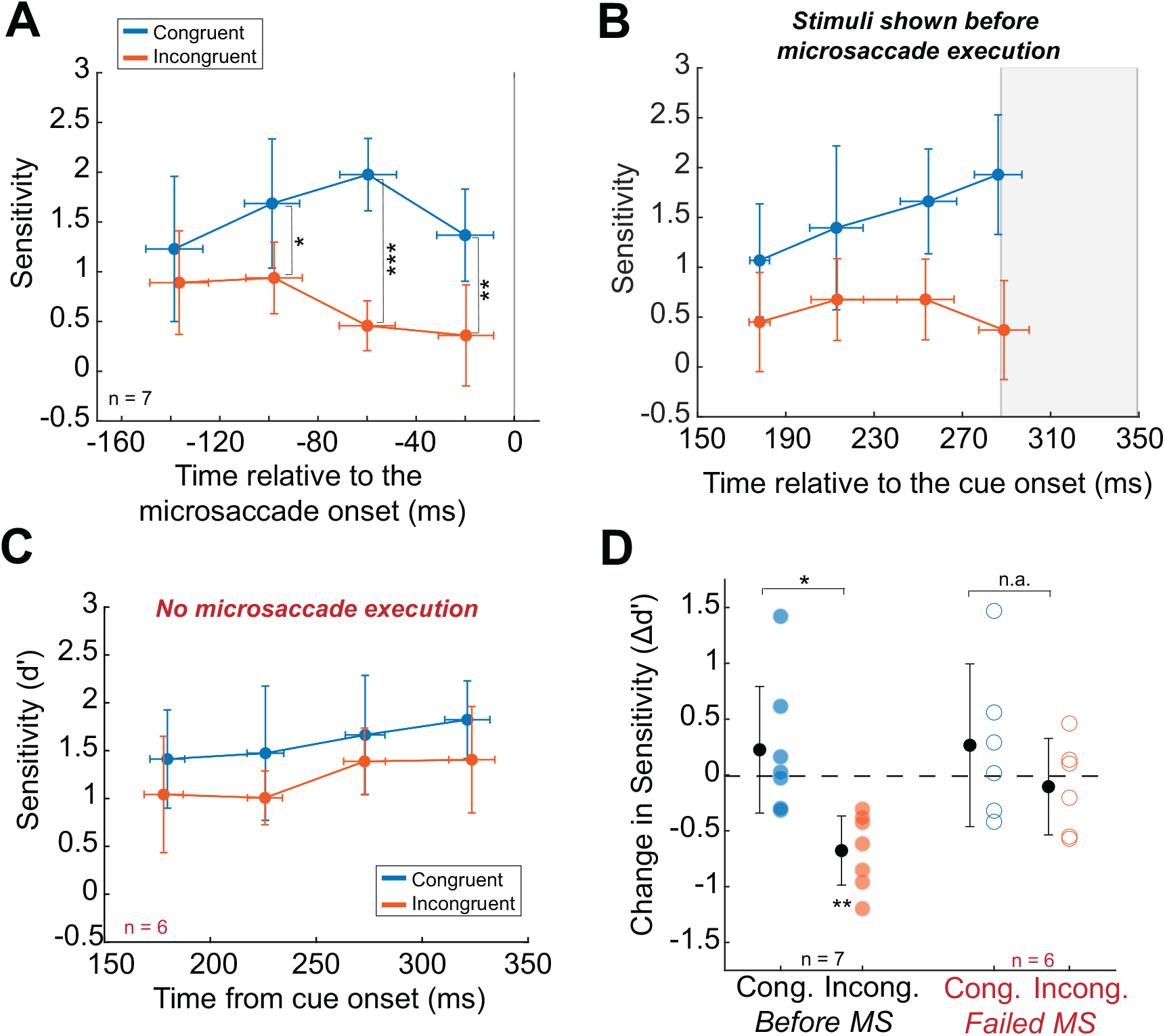
Assessing the influence of covert attention. Average performance as a function of time relative to microsaccade (A) and relative to the cue (B) for congruent (blue) and incongruent (orange) trials. Each non-overlapping time bin in A and B covers 40 milliseconds. Shaded gray region indicates 95% within-subject confidence interval of microsaccade onset time. C) Average performance relative to the cue onset in congruent and incongruent trials in which observers failed to perform microsaccades. Each non-overlapping time bin in C covers 50 milliseconds. In A-C, ^*^*p<*0.05, ^**^*p<*0.01 (two-way repeated-measures ANOVA (factors: condition x time) with Tukey-Kramer corrected *t*-tests; comparisons between congruent and incongruent at each time bin are shown). D) Same data shown in B and C averaged over time within each group. ^*^*p<*0.05, ^**^*p<*0.01 (paired two-sided *t*-tests with Bonferroni corrected p-values). Vertical error bars in graphs are 95% within-subject confidence intervals and horizontal error bars represent standard deviations.

To rule out the possible influence of covert attention, we first examined changes in performance with respect to the cue onset rather than the microsaccade onset. To this end, we focused only on the trials with stimulus presentations occurring before the microsaccade onset. This was to ensure that factors such as saccadic suppression and differences in stimulus eccentricity did not influence performance. Therefore, the only influence on performance could be attentional factors either related to the microsaccade preparation itself or endogenous cueing of attention. As shown in Figure 4B, performance in congruent trials was higher than the performance in the incongruent trials, however, this pattern did not interact with time relative to the cue onset (two-way repeated-measures ANOVA, condition:*F*(1,6)= 31.39, *p*= 0.0014; time:*F*(3,18)= 1.470, *p*_GG_= 0.273; interaction:*F*(3,18)= 3.15, *p*_GG_= 0.09). This is in contrast with the pattern of performance locked in time to the onset of the microsaccade, where significant differences between congruent and incogruent trials were found to vary over time. Performance in congruent and incongruent trials did not show signifianct temporal modulations when locked to the time of cue onset, whereas a more obvious and significant interaction was found when the same data were locked relative to the microsaccade onset.

### Visual Discrimination in Failed Microsaccades Trials

To further rule out any contribution of attention unrelated to saccadic factors, we examined the trials in which subjects failed to perform microsaccades, *i*.e., trials in which a directional motor cue was present but subjects did not perform a microsaccades. On average the 7 subjects who completed data collection failed to execute microsaccades in 28% ± 14% of the trials with a directional cue. Failed microsaccade trials from 5 subjects were analyzed along with 1 additional observer who contributed only failed trials, for a total of 6 subjects with failed microsaccade trials. 2 subjects were excluded from this analysis due to too few failed microsaccade trials. If voluntary shifts of covert attention contributed to the impairment in performance during the incongruent trials, the same effects should be present in trials when subjects fail to perform a microsaccade following the central cue. However, no difference in performance between congruent and incogruent trials and no modulation in performance over time was found following the cue onset as shown in Figure 4C (two-way repeated-measures ANOVA, condition:*F*(1,5)= 2.396, *p*= 0.1823; time:*F*(3,15)= 3.069, *p*_GG_= 0.0727; interaction:*F*(3,15)= 0.119, *p*_GG_= 0.884)). In fact, as illustrated in Figure 4D, in the initial 350 milliseconds following the onset of the central cue, there is no difference in performance between the congruent and incongruent trials when observers failed to perform microsaccades (Bonferroni corrected t-test, two-sided, paired: *p*= 0.3764, *BF*_10_= 0.8042) nor an impairment compared to baseline (Bonferroni corrected t-test, two-sided, paired: *p*= 1.000, *BF*_10_= 0.414). On the contrary, when the target was presented before microsaccade execution, performance in the incongruent trials was impaired relative to baseline (Bonferroni corrected t-test, two-sided, paired: *p*= 0.0035, *BF*_10_= 26.278) and the congruent trials (Bonferroni corrected t-test, two-sided, paired: *p*= 0.0102, *BF*_10_= 11.286).

These results indicate that the modulations in fine spatial vision preceding microsaccade onset are linked to the execution of the microsaccade, rather than being an effect of cueing or voluntary attention independent of eye movements. Observing a coherent time course relative to the cue could reveal a cue-locked visual mechanism that does not depend on the dynamics of saccade-related modulations. No such modulations in perceptual performance was present when observers fail to execute a microsaccade.

## Discussion

Microsaccades are often regarded as a marker of extrafoveal covert shifts of attention; they are generally oriented toward covertly attended peripheral locations following a particular time course (***Engbert and Kliegl, 2003***; ***Hafed and Clark, 2002***; ***Tian et al., 2016***; ***Liu et al., 2022, 2024***; ***Brandolani et al., 2025***), and have been associated with perceptual enhancements at those locations (***Hafed et al., 2011***; ***Hafed, 2013***; ***Yuval-Greenberg et al., 2014***; ***Chen et al., 2015***). On the other hand, the effects of microsaccades at the foveal scale in the presence of more complex foveal stimuli have been overlooked. At the foveal scale microsaccades are known to precisely redirect the preferred locus of fixation (***Ko et al., 2010***; ***Poletti et al., 2013***), and play a crucial role in shaping foveal vision as they actively contribute to visual exploration of complex foveal stimuli (***Shelchkova et al., 2019***). Further, microsaccades are characterized by a tight interplay with selective attention, not only at the larger scale (***Yuval-Greenberg et al., 2014***; ***Chen et al., 2015***; ***Hafed et al., 2015***), but also within the central 1-degree fovea (***Shelchkova and Poletti, 2020***; ***Guzhang et al., 2024***). In this study, we examined the temporal dynamics of perceptual changes in foveal vision around the time of microsaccades. To this end, we briefly presented fine spatial stimuli at variable times relative to the onset of an instructed microsaccade. Our findings show how foveal vision is dynamically re-shaped with each microsaccade. Perception is first prioritized at the target vs non target location right before the microsaccade onset; this perceptual redistribution of sensitivity across the foveola peaks ≈70 milliseconds before microsaccade onset, and is strongly mediated by a perceptual impairment at the location isoeccentric to the microsaccade target. Further, the rapid change in sensitivity across the foveola is associated with microsaccades rather than endogenous shifts of covert attention occurring independently of microsaccade preparation. Consistent with previous work (***Zuber and Stark, 1966***; ***Hafed and Krauzlis, 2010***; ***Intoy et al., 2021***), during microsaccade execution, fine visual discrimination is substantially impaired. Then, within ≈100 milliseconds upon microsaccade landing performance at the target location rapidly improves reaching levels comparable to those achieved during the pre-microsaccadic enhancement period.

The findings reported here align with recent neurophysiological evidence showing that, just before a microsaccade toward a small visual stimulus, neural activity in macaque V1 is enhanced at both the current stimulus location and the anticipated location of the stimulus post-microsaccade (***Bouhnik et al., 2025***). This enhancement is accompanied by increased neural synchronization at the target site, suggesting the involvement of a premotor attentional signal that prepares the visual system for the stimulus expected to fall on the preferred fixation locus after the microsaccade lands (***Melloni et al., 2009***; ***Bouhnik et al., 2025***). Notably, the peak in microsaccade-related neural synchronization within striate and extrastriate cortex occurs approximately 100 milliseconds after microsaccade onset (***Leopold and Logothetis, 1998***; ***Meirovithz et al., 2012***; ***Lowet et al., 2018***; ***Nativ et al., 2025***), a time point that coincides with the post-saccadic recovery of visual performance observed in our study. This temporal correspondence suggests that the post-microsaccadic synchronization may facilitate the rapid processing of visual input following saccade landing.

The presence of highly localized perceptual enhancements that follow a systematic temporal course around the time of microsaccades in the central fovea raises important questions about their relationship to microsaccade-related effects in the visual periphery. Interestingly, foveal enhancements have typically been studied in isolation, without concurrent extrafoveal stimulation, while peripheral effects have mostly been examined under conditions in which subjects fixate on an impoverished foveal stimulus. Yet, under natural viewing conditions, both foveal and extrafoveal inputs are continuously present and explored in a continuous fixation–saccade cycle. Do the perceptual modulations associated with microsaccades in the fovea and periphery coexist in natural viewing conditions, and how do they interact? Does a microsaccade targeting a detail within the foveola elicit perceptual changes not only at its goal location but also peripherally? Understanding the timing and spatial dynamics of these pre-microsaccadic attentional modulations in naturalistic settings will require further investigation, particularly in contexts where both foveal and peripheral stimuli are task relevant.

Extensive research has characterized the perceptual benefits preceding large saccades near their goal locations (***Kowler et al., 1995***; ***Hoffman and Subramaniam, 1995***; ***Deubel and Schneider, 1996***; ***Gersch et al., 2004***; ***Rolfs et al., 2011***; ***Rolfs and Carrasco, 2012***; ***Harrison et al., 2013***; ***Li et al., 2016***; ***Ohl et al., 2017***; ***Kroell and Rolfs, 2021***). Among these, some behavioral studies investigating the temporal dynamics of this phenomenon consistently report the onset of perceptual modulations beginning 50-100 milliseconds and peaking 25 milliseconds before the saccade begins. For example, ***Rolfs and Carrasco (2012***) observed improved discrimination and perceived contrast emerging ≈100 milliseconds before saccade onset, and peaking ≈25 milliseconds before the onset. Similarly, ***Ohl et al. (2017***) reported a gain in perceptual orientation tuning in the 100 milliseconds preceding the saccade onset and a narrowing of the orientation tuning immediately before the saccade onset. Our findings reveal that pre-microsaccadic perceptual enhancements also begin ≈100 milliseconds before its onset, consistent with the temporal onset of pre-saccadic enhancements. However, in our study, the difference in performance between baseline and congruent trials peaked within a broader window that extended for 79 ± 48 milliseconds before the microsaccade onset, with the peak time of the enhancement occurring earlier than previously reported for larger saccades (average peak time: 70 ± 23 milliseconds before onset). This discrepancy in the time course of pre-saccade and pre-microsaccade effects could either be due to variations in experimental design and stimuli or to differences in the temporal dynamics of the mechanisms underlying the control of smaller and larger saccades. Microsaccades are, in fact, characterized by significantly longer reaction times compared to saccades (***Kalesnykas and Hallett, 1994***; ***Shelchkova et al., 2019***; ***Shelchkova and Poletti, 2020***) hence, similar differences may be reflected in the dynamics of premicrosaccadic effects.

Presaccadic perceptual enhancement is known to be feature-specific, often favoring higher spatial frequencies and sharpened orientation tuning at the saccade target (***Li et al., 2016, 2019***; ***Ohl et al., 2017***). Some studies showed that, just before saccades to targets at 10^◦^ eccentricity, perception of 1.5-2.0 cpd gratings improved relative to 1.0 cpd gratings (***Li et al., 2016, 2019***). More recent work has extended these findings to a broader frequency range and found that presaccadic enhancement narrows the sensitivity bandwidth, increases peak sensitivity, and shifts preferred spatial frequency upward—up to 1.6 cpd for horizontal targets (***Kroell and Rolfs, 2021***). However, frequencies above ≈2.5 cpd received less benefit, suggesting an upper limit to this enhancement. Recent studies have shown that these effects vary across the visual field. For example, when saccade targets were closer to fixation (6^◦^ eccentricity) and tested with a wider frequency range (0.5–10 cpd), peak enhancement shifted to ≈5 cpd and was greater for horizontal than vertical saccades, revealing a dependency on target location, saccade direction, and visual field meridian (***Hanning et al., 2024***; ***Kwak et al., 2024***). In the present study, we used bar stimuli containing a relatively broad range of spatial frequencies that could be used to perform a coarse discrimination task. Although these stimuli allowed us to restrict stimulation to a small area at the saccade target, they did not clarify whether the observed perceptual modulations are narrowly tuned or broadly distributed across spatial frequencies. As the fovea supports perception across a broader range of spatial frequencies than the visual periphery, and it is primarily used to resolve fine detail, microsaccade-related enhancement may primarily affect higher spatial frequencies.

Our findings demonstrate that microsaccades are accompanied by a temporally structured sequence of spatially selective modulations in foveal vision. Fine visual discrimination is prioritized at the microsaccade target during the ≈100 milliseconds preceding movement onset, is transiently suppressed during the movement execution, and rapidly recovers within ≈100 milliseconds after the stimulus is brought to the preferred locus of fixation. Although microsaccades are known for their high accuracy and precision, shifting the preferred retinal locus by fractions of a degree to compensate for non-uniform foveal sensitivity, our results show that their perceptual benefits begin well before the microsaccade begins. These findings reveal that microsaccades do more than precisely redirect gaze: they dynamically reshape foveal processing over time, supporting the fine-grained selection of visual information during active fixation.

## Methods and Materials

### Observers and Apparatus

Data were collected from 9 observers (1 author and 8 naïve; age 20–30 years; 7 females, 2 males). One na ive observer withdrew before completing the study. Data presented here are from 8 observeres (7 female, 1 male). All observers performed a visual acuity screening using a Snellen Chart prior to participation to ensure binocular 20/20 acuity. Naive observers were paid 15 USD dollars per hour. Ethical approval for this research study was obtained from the University of Rochester’s Research Subjects Review Board. Observers attended an initial screening session, which involved a comprehensive explanation of the experiment and a thorough review of the consent form materials. After understanding the information and voluntarily agreeing to participate, informed consent was obtained and documented. Observers viewed stimuli displayed on an LCD monitor (ASUS PG259QN), shown with a vertical refresh rate of 200 Hz and a spatial resolution of 1920 X 1080 pixels. The monitor was 1550 mm away from the observer (1 pixel = 0.62’). Vertical and horizontal eye movements were recorded with high precision by means of a custom-made digital DPI eye tracker with a sampling rate of 340 Hz. The head was immobilized by means of a dental-imprint bite bar and head holder to reduce noise in the eye-tracking signal.

### Experimental Design

Data were collected over multiple experimental sessions. Each observer completed 11-18 sessions (total trials discarded across observers: 48% ± 11%) with each session lasting approximately 1 hour. All sessions required a preliminary setup that included adjusting the observer in the apparatus, tuning the eye tracker for optimal performance, and executing a two-step gaze-contingent calibration procedure. In the first step of the calibration procedure, referred to as automatic calibration, observers sequentially fixated on each of the nine points of a 3 x 3 grid for about 2 seconds, as in customary eye tracking experiments. These nine points were located 150 pixels apart on the horizontal and vertical axes. In the second phase, referred to as manual calibration, observers confirmed or refined the mapping given by the automatic calibration by fixating on each of the nine points of the grid while the location of the line of sight estimated by the previous calibration step was displayed in real-time on the screen. Observers used a joypad to correct the estimated gaze location of the screen if necessary. These corrections were then incorporated into the camera pixel-to-monitor pixel transformation. This dual-step calibration procedure allows for a more accurate 2D localization of the line of sight by approximately one order of magnitude (***Poletti et al., 2013***; ***Poletti and Rucci, 2016***). The manual calibration was repeated for the central position after each trial to compensate for head movements that may occur during data collection.

During the task, observers were instructed to fixate on a tiny 3’ x 3’ central marker located between two 5’ x 5’ boxes shown at the center of the display. Observers maintained fixation for 400-600 milliseconds until a cue appeared at the center of the screen. In motor cue trials Figure 1B, the central cue was a horizontal line pointing either to the left or right location with equal probability. Observers were instructed to shift their gaze as quickly as possible toward the location indicated by the cue. In the neutral cue trials Figure 1B, the central cue was a fixation cross. This cue instructed observers to maintain fixation on the central marker throughout the rest of the trial. After a variable delay (up to 600 milliseconds) from the cue offset, high-acuity stimuli, 7’ x 3’ bars tilted 45^◦^ to the left or right of vertical, were presented (50 milliseconds) simultaneously at both box locations. The orientation of each stimulus was randomized. 500 milliseconds after stimuli onset a response cue appeared instructing observers to report the orientation of the line shown at the cued location. Observers reported their response by pressing a button on the joypad. The cue type (neutral or motor) and the location of the response cue were randomly and independently determined on each trial.

### Data Analysis

#### Eye Movement Processing

Recorded oculomotor traces were examined offline to determine the precise timing of stimulus presentation relative to the executed gaze shifts. Only blink-free trials with optimal tracking of the first and fourth Purkinje images were selected for data analysis. Classification of eye movements, such as microsaccades, saccades, and drift, were performed automatically based on eye velocity and then validated for each observer by trained laboratory personnel with extensive experience in classifying eye movements. Eye movement traces were filtered with a Savitzky-Golay filter before calculating instantaneous speed. Data segments in which the eye moved by more than 5’ reaching an instantanous speed of 3.5°/s were marked as possible saccades, and their onset and offset were defined as the initial and final times at which the eye speed exceeded and returned to below 2^°^/s, respectively. Saccades with an amplitude of less than 0.5^°^ (30’) were defined as microsaccades. Consecutive events closer than 15 milliseconds were merged together. Periods that were not classified as saccades or blinks were labeled as drifts. The average eye position along the horizontal axis and the average instantaneous eye speed relative to the time of microsaccade onset are shown in Figure 5.

#### Trial Selection Criteria

Trials in which the target was presented before the microsaccade onset were discarded if the eye position was outside of a 10’ region surrounding the fixation marker during stimulus presentation. *Microsaccade Trials:* Only trials in which microsaccades occurred within 475 milliseconds of the motor cue were analyzed as microsaccade trials shown in Figure 2B. A small portion of the data was excluded for microsaccade latencies greater than 475 milliseconds (18% ± 18%, where only 2 out of the 7 subjects had rates larger than 13%). Each microsaccade trial was classified by the time of the stimuli offset relative to the onset of the microsaccade. In the “before” trials the target offset must occur before the microsaccade onset. In the “during” trials the target must be presented during the peak speed of the microsaccade. In the “after” trials the target onset must occur after the microsaccade offset. Trials were discarded if microsaccade landing position was greater than 14’ on the vertical axis and greater than 10’ on the horizontal axis from the target location. *Neutral Trials:* Only neutral trials in which the observer maintained fixation (without microsaccades) for the entire trial were analyzed. *Failed Microsaccade Trials:* Trials in which the observer’s maintained fixation despite being presented with the microsaccade motor cue were analyzed as failed microsaccade trials.

#### Performance Analysis

*Smoothing Performance Relative to Microsaccade Onset:* To calculate performance over time, trials were grouped based on the stimulus offset time relative to the microsaccade onset time for both congruent and incongruent conditions, covering the interval from -175 milliseconds to 300 milliseconds. Performance (evaluated as percent of correct responses) was averaged within a 50 millisecond sliding window, advancing in 1 ms steps (with 24 ms overlap). Task performance was also evaluated as d-prime. A “right” response to a right tilted stimulus was classified as a Hit, whereas a “right” response to a leftward stimulus was classified as a False Alarm. The rate of hits and false alarms were z-score transformed to calculate the dprime. *Calculating Baseline Performance in Neutral Trials:* Because performance in Neutral trials varied slightly over time relative to cue onset, we matched microsaccade and no-microsaccade trial performance based on the timing of stimulus onset with respect to the cue. Accordingly, in Figures 3C and 4A, baseline performance for each microsaccade category (*i*.*e*., before, during, and after) was estimated using neutral trials whose delay times (the interval between cue onset and target onset) fell within one standard deviation of the mean delay time for the corresponding microsaccade category. *Pre-Microsaccadic Enhancement Analysis:* To determine the onset and offset of the pre-microsaccadic enhancement in congruent trials, we subtracted performance in neutral, delay-matched trials from performance in microsaccade-congruent trials. The onset of the perceptual enhancement was defined as the time point at which this performance difference exceeded 5% for at least 15 ms. The offset was defined as the time when the performance difference dropped back below this threshold following the onset.

#### Statistical Analysis

Statistical analyses were performed in MATLAB using the Statistics and Machine Learning Toolbox. Data were analyzed using one-way and two-way repeated-measures ANOVA. When Mauchly’s test indicated a violation of the sphericity assumption, *p*-values were corrected using the Green-house–Geisser method and are reported as *p*_GG_. Pairwise comparisons were adjusted for multiple testing using either Bonferroni or Tukey–Kramer corrections, as appropriate. Bonferroni correction was applied to planned pairwise comparisons that were not part of the omnibus ANOVA, whereas Tukey–Kramer adjustment was used for pairwise comparisons included in the omnibus ANOVA. Bayes factors were computed using paired Bayesian two-sided *t*-tests with Jeffreys–Zellner-Siow (JZS) priors, implemented via the *BayesFactor* Toolbox (https://zenodo.org/badge/latestdoi/162604707) using MATLAB, (RRID:SCR_001622). Effect sizes are reported as Cohen’s *d*. 95% within-subject confidence intervals were calculated using the inverse of Student’s *t* cumulative distribution function and are shown for each sample mean in the figures. We estimated the required sample size based on effect sizes reported in prior studies of pre-microsaccadic attention using similar paradigms (***Shelchkova and Poletti, 2020***; ***Guzhang et al., 2024***), which typically observed large effect sizes. A power analysis assuming comparable size effects with an average within-subject correlation of *ρ*= 0.5 across conditions yielded a sample size of 7.

## Acknowledgments

We are grateful to current and former members of the Active Perception Lab for their feedback during this project. This work would not have been possible without support from NIH R01 EY029788-01 and NIH EY001319 grant to the Center for Visual Science at the University of Rochester.

## Supplemental Figures

**Figure 5. Supplementary 1:**
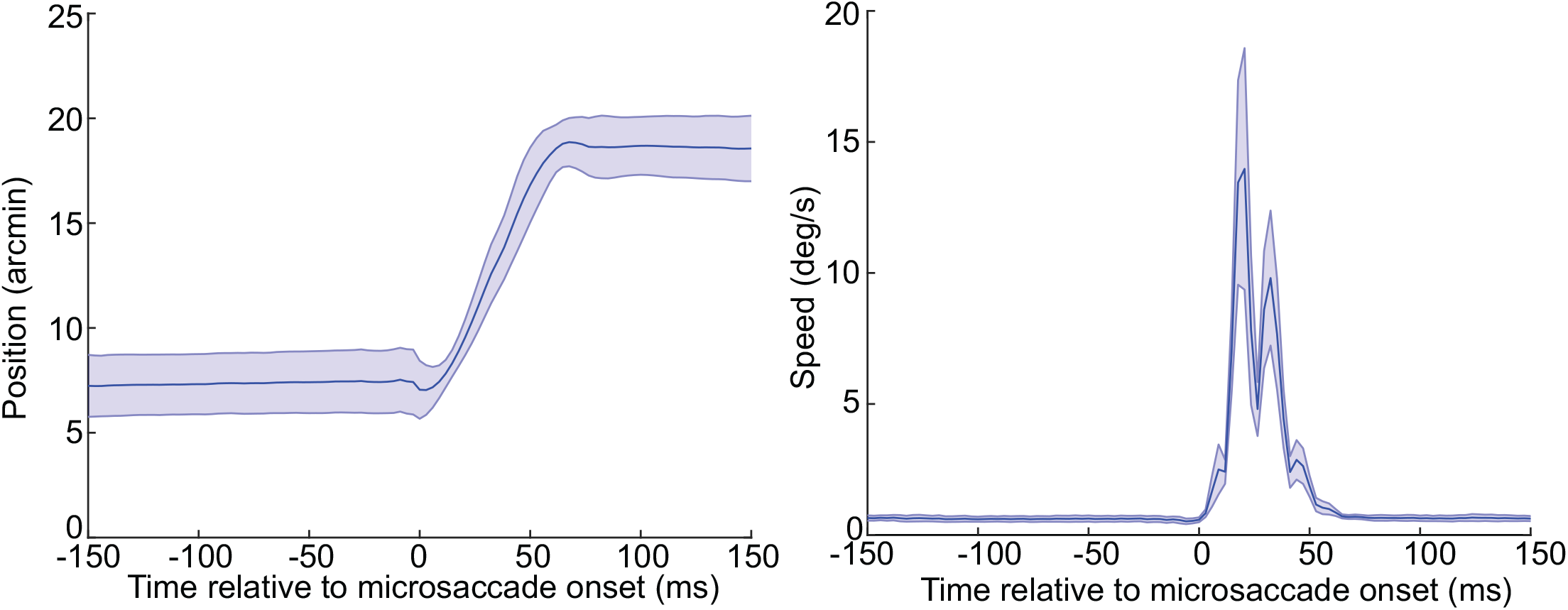
Average horizontal gaze position and two-dimensional eye speed relative to microsaccade onset. Shaded error bars are standard deviation across individual subjects.

**Figure 6. Supplementary 2:**
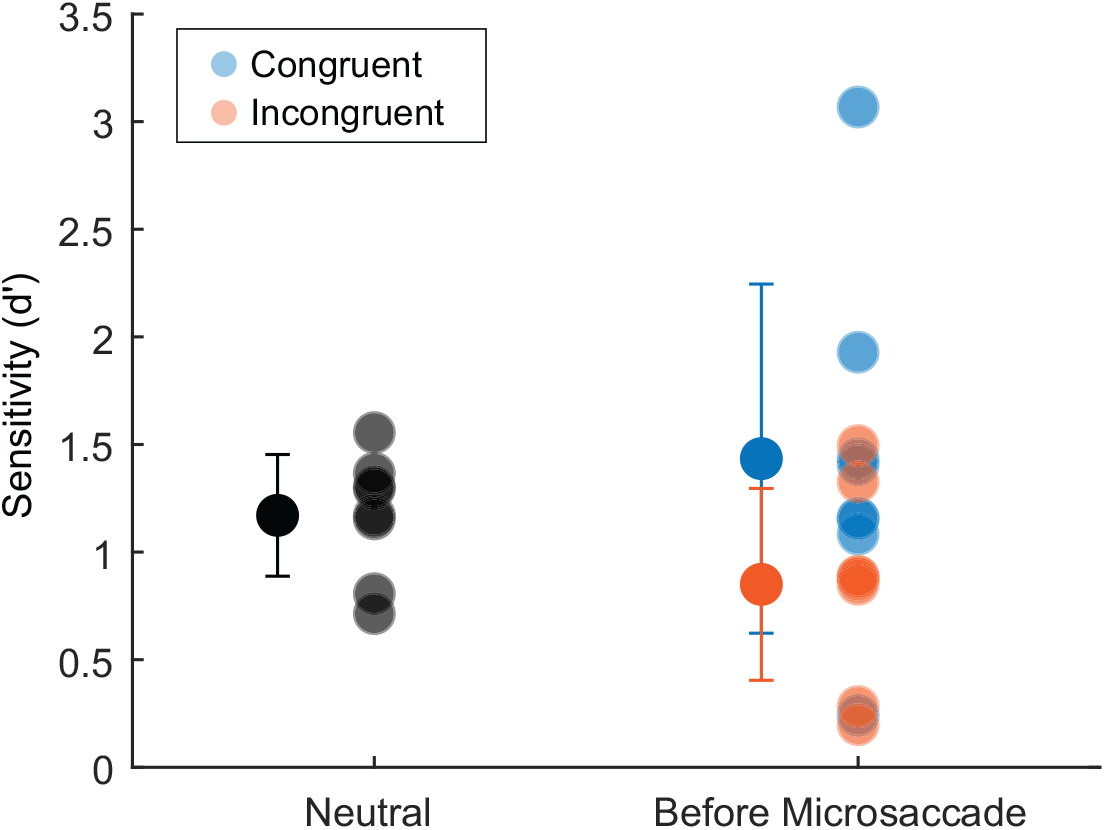
Average performance in congruent (blue) and incongruent (orange) trials -150 to -100 ms before the microsaccade onset compared to delay-matched neutral trials.

